# PhenotypeToGeneDownloaderR: automated multi-source retrieval and validation of phenotype-associated genes

**DOI:** 10.64898/2026.05.02.721360

**Authors:** Muhammad Muneeb, David B. Ascher

## Abstract

**Motivation:** Identifying phenotype-associated genes is a common first step in polygenic risk score construction, enrichment testing, target prioritisation and variant interpretation, but relevant evidence is distributed across heterogeneous databases with different interfaces, formats and evidence models.

**Results:** We present PhenotypeToGeneDownloaderR, a phenotype-guided R/Python pipeline for automated gene retrieval, harmonisation, symbol validation and cross-source summary analysis. Given a phenotype term, the pipeline queries integrated biological databases, standardises per-source outputs, combines gene lists, validates retrieved symbols against the NCBI human gene reference and generates summary tables and visualisations. Across 13 clinically relevant phenotypes and 13 databases, PhenotypeToGeneDownloaderR generated 136,487 raw gene retrievals, with at least one source returning genes for every phenotype. Across all 13 phenotypes, 100,175 of 114,345 combined input symbols were retained after direct or synonym-based validation, corresponding to an 87.6% validation rate. Cross-source overlap was low, supporting the complementarity of integrated evidence sources. Against an HPO/ClinVar/OMIM-derived gold standard, the pipeline recovered 1,039 of 1,056 known phenotype-associated genes, corresponding to 98.4% recall. PhenotypeToGeneDownloaderR provides a lightweight, reproducible upstream framework for generating candidate gene sets for downstream prioritisation and interpretation.

**Availability and implementation:** PhenotypeToGeneDownloaderR is implemented in R and Python, released under the MIT licence, and available at https://github.com/MuhammadMuneeb007/PhenotypeToGeneDownloaderR.

**Supplementary information:** Supplementary data are available online.

## Introduction

Identifying genes associated with a phenotype or disease is a frequent starting point for human genetics and bioinformatics analyses, including enrichment testing, polygenic risk score construction, target prioritisation and variant interpretation [1, 2, 3]. However, phenotype–gene evidence is distributed across resources that differ in scope, interface, terminology, evidence type and output format. Clinical resources such as ClinVar [4], OMIM [5] and HPO [6] capture curated disease and phenotype associations; pathway and protein resources such as KEGG [7], Reactome [8], STRING [9] and UniProt [10] capture functional or interaction evidence; and integrative resources such as Open Targets [11], DisGeNET [12] and the GWAS Catalog [13] provide disease, genetic association and multi-evidence links. As a result, researchers often need to query multiple resources independently and manually reconcile heterogeneous gene identifiers before downstream analysis.

Existing packages and web resources are valuable, but many focus on a single database, require pre-defined gene sets, or emphasise downstream enrichment rather than phenotype-first retrieval and harmonisation [14, 15, 16]. This creates a practical bottleneck: the same phenotype may yield different candidate gene sets depending on which source is queried, how terms are matched and how gene symbols are reconciled.

PhenotypeToGeneDownloaderR addresses this gap by providing a unified phenotype-first workflow for multi-source gene retrieval, standardisation and cross-source evidence aggregation. Given a phenotype term, the pipeline retrieves candidate genes from integrated biological databases, writes standardised per-source outputs, combines gene lists, validates symbols against the NCBI human gene reference and produces reproducible summary outputs for downstream prioritisation. The tool is intended as an upstream triage layer: it generates analysis-ready candidate gene sets and source-support summaries, rather than claiming to establish causal gene–phenotype relationships.

## Materials and Methods

PhenotypeToGeneDownloaderR consists of two linked components: an R-based retrieval layer and a Python-based downstream analysis layer. Given a phenotype term, the retrieval layer queries integrated biological databases and writes standardised per-source CSV outputs. The downstream layer combines cross-source results, summarises recovery patterns, and generates analysis-ready outputs. Retrieved gene symbols are additionally cross-referenced against the NCBI human gene reference to distinguish valid symbols from artefacts and to resolve synonyms to current official symbols.

For the present benchmark, the pipeline was evaluated across 13 clinically relevant phenotypes and 13 integrated biological databases spanning literature, clinical genetics, ontology, pathway, expression, protein-function, and integrative evidence types. The overall workflow is shown in Figure 1.

**Fig. 1.**
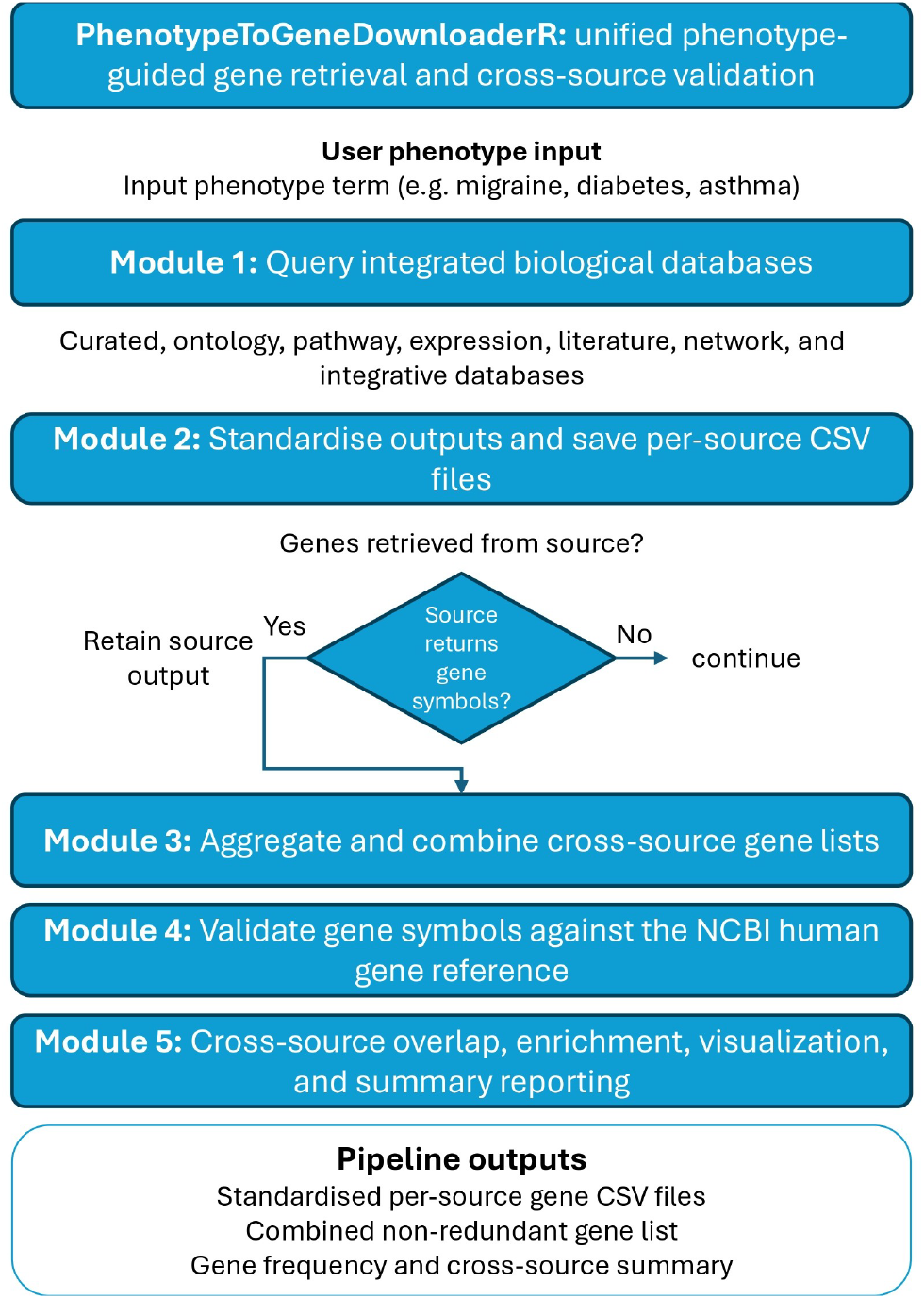
Overview of the PhenotypeToGeneDownloaderR workflow. A phenotype term is used to query integrated biological databases, generate standardised per-source CSV outputs, combine cross-source gene lists, validate gene symbols against the NCBI human gene reference, and produce downstream summary analyses and visualisations.

## Results and Discussion

Across 13 clinically relevant phenotypes and 13 integrated biological databases, PhenotypeToGeneDownloaderR generated 136,487 raw gene retrievals, with at least one source returning genes for every phenotype. Twelve of 13 databases returned genes for at least one phenotype, while OMIM and GTEx achieved complete phenotype coverage in this benchmark. Open Targets and the GWAS Catalog contributed the largest total gene yields, whereas lower-yield resources such as Reactome, KEGG and Gene Ontology contributed more selectively, reflecting differences in database scope, terminology, query behaviour and evidence model (Table 1; Supplementary Section S1).

**Table 1.**
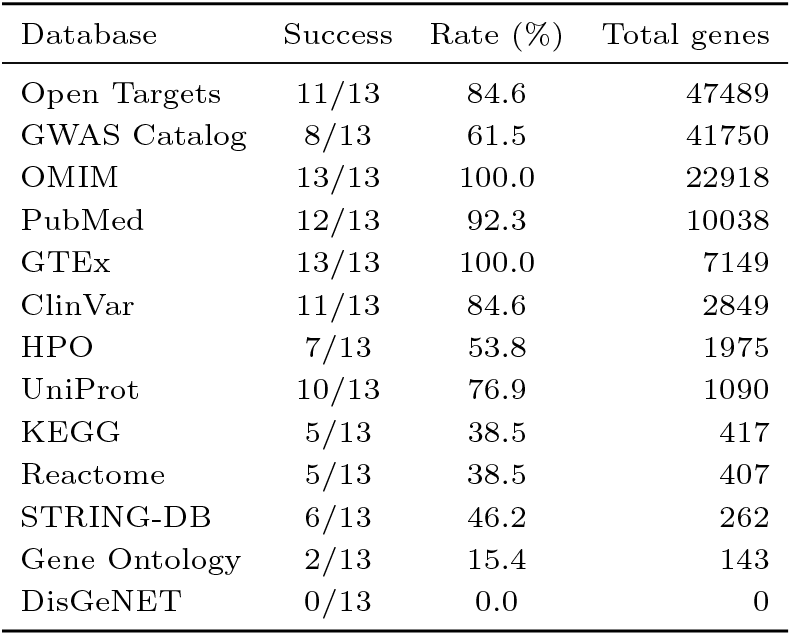
Compact summary of per-database retrieval performance across 13 phenotypes. Success indicates the number of phenotypes for which a database returned at least one gene symbol. Total genes indicates the sum of unique genes returned across all phenotypes prior to cross-source merging and symbol validation.

Gene-symbol validation and harmonisation retained 100,175 of 114,345 combined input symbols across all 13 phenotypes, corresponding to an overall validation rate of 87.6%. This step removed non-gene artefacts and unresolved identifiers while rescuing outdated or alternative aliases through synonym-based mapping to current official symbols. In total, 4,912 validated symbols were resolved through synonym mapping, representing 4.9% of the validated set (Supplementary Section S2). These results show that the pipeline can convert heterogeneous source-level outputs into cleaner candidate gene sets suitable for downstream review and interpretation.

KEGG and DisGeNET require cautious interpretation in the current benchmark. KEGG returned source-level records for some phenotypes, but these produced zero validated genes after the current parsing and symbol-validation step, indicating an implementation-level parsing limitation. DisGeNET returned no genes across the evaluated phenotypes because API access was required but was not available during these runs. These results are therefore reported as access or implementation limitations rather than evidence that these resources lack relevant phenotype–gene associations.

Cross-source overlap was low overall, supporting the complementarity of the integrated evidence sources. Most genes were supported by a single database, while a smaller subset was recovered by multiple independent sources and may represent higher-confidence candidates for downstream prioritisation. GWAS Catalog, GTEx, Open Targets and OMIM contributed substantial unique gene fractions, indicating that each source captures partly distinct biological or evidence dimensions (Supplementary Section S3).

To assess recovery of established phenotype-associated genes, we compared the retrieved gene sets against an HPO/ClinVar/OMIM-derived gold standard. PhenotypeToGeneDownloaderR recovered 1,039 of 1,056 curated known phenotype-associated genes, corresponding to 98.4% recall. Precision@20 analysis further supported the ability of the source-frequency ranking to prioritise known phenotype-associated genes among the highest-ranked candidates, with a mean Precision@20 of 64.1% across phenotypes with non-empty gold-standard sets (Supplementary Section S4). Downstream enrichment using g:Profiler produced significant enrichment results for all 13 phenotypes, supporting the biological coherence of the generated candidate gene sets while remaining a downstream characterisation rather than causal validation (Supplementary Section S5).

The benchmark also showed practical usability. Total runtime across 13 phenotypes was 94.1 minutes, with a mean runtime of 7.2 minutes per phenotype and modest peak memory use. Empirical comparison with implemented modules showed that the full pipeline recovered a larger fraction of gold-standard genes than the GWAS Catalog-only or Open Targets-only modules, while the top 500 source-ranked genes submitted to g:Profiler retained 793 of 1,056 gold-standard genes (75.1%) (Supplementary Section S6). Together, these results support the use of PhenotypeToGeneDownloaderR as a lightweight and reproducible first-pass framework for phenotype-guided candidate gene retrieval and cross-source evidence aggregation. The tool is not designed to prove causal gene–phenotype relationships directly; rather, it provides harmonised, source-supported candidate gene sets for downstream interpretation, enrichment analysis and expert review.

## Supporting information

Supplemental Material All

## Availability and implementation

PhenotypeToGeneDownloaderR is freely available as an open-source software pipeline at https://github.com/MuhammadMuneeb007/PhenotypeToGeneDownloaderR. The repository is released under the MIT licence and includes the complete R-based phenotype-to-gene retrieval scripts, Python-based downstream analysis scripts, dependency files, documentation and example usage instructions. The current release described in this manuscript is version 1.0.0. The retrieval layer is implemented in R and coordinated through download genes.R, which executes source-specific database modules for phenotype-guided gene retrieval. The downstream analysis layer is implemented in Python and includes scripts for source coverage analysis, gene-symbol validation, cross-source overlap assessment, known-gene recovery, enrichment analysis and summary visualisation. Installation requirements are provided through requirements.R, requirements.txt and environment.yml, enabling users to reproduce the R and Python environments required for execution. Given a phenotype term, the pipeline generates per-source output files, combined cross-source gene lists, source-summary tables, validation outputs, overlap statistics and publication-ready summary plots. Example commands, expected output structure and documentation are provided in the GitHub repository. The benchmark phenotypes and supplementary outputs described in this manuscript provide test cases for reproducing retrieval coverage, symbol validation, cross-source complementarity, known-gene recovery and runtime analyses. PhenotypeToGeneDownloaderR is intended as a reproducible upstream candidate-gene retrieval and prioritisation framework for downstream analyses such as enrichment testing, polygenic risk score development, target prioritisation and variant interpretation. It does not infer causal gene–phenotype relationships directly; instead, it harmonises and summarises evidence from multiple public biological databases to support downstream review and interpretation.

## Author contributions statement

M.M. wrote the first draft of the manuscript and wrote, tested, and documented the code. M.M. analysed the results. D.A. reviewed and edited the manuscript. All authors contributed to the methodology.

## Data availability

The source code, documentation, example commands and example output files are available in the GitHub repository at https://github.com/MuhammadMuneeb007/PhenotypeToGeneDownloaderR. The underlying databases are publicly accessible: PubMed (https://pubmed.ncbi.nlm.nih.gov/), OMIM (https://www.omim.org/), ClinVar (https://www.ncbi.nlm.nih.gov/clinvar/), HPO (https://hpo.jax.org/), Gene Ontology (https://geneontology.org/), KEGG (https://www.kegg.jp/), Reactome (https://reactome.org/), STRING (https://string-db.org/), GTEx (https://gtexportal.org/), UniProt (https://www.uniprot.org/), Open Targets (https://www.opentargets.org/), DisGeNET (https://www.disgenet.org/), and the GWAS Catalog (https://www.ebi.ac.uk/gwas/).

## Acknowledgments

Not applicable.

